# Novel cell-to-cell interactions revealed by cryotomography of a DPANN coculture system

**DOI:** 10.1101/2024.05.20.594898

**Authors:** Matthew D Johnson, Doulin C Shepherd, Hiroyuki D. Sakai, Manasi Mudaliyar, Arun Prasad Pandurangan, Francesca L Short, Paul D. Veith, Nichollas E Scott, Norio Kurosawa, Debnath Ghosal

## Abstract

DPANN is a widespread and highly diverse group of archaea characterised by their small size, reduced genome, limited metabolic pathways, and symbiotic existence. Known DPANN species are predominantly obligate ectosymbionts that depend on their host for their survival and proliferation. Despite the recent expansion in this clade, the structural and molecular details of host recognition, host-DPANN intercellular communication, and host adaptation in response to DPANN attachment remain unknown. Here, we used electron cryotomography (cryo-ET) to reveal that the *Candidatus* Micrarchaeota (ARM-1) interacts with its host, *Metallosphaera javensis* through intercellular proteinaceous nanotubes. These tubes (∼4.5 nm wide) originate in the host, extend all the way to the DPANN cytoplasm and act like tunnels for intercellular exchange. Combining cryo-ET and sub-tomogram averaging, we revealed the *in situ* architectures of host and DPANN S-layers and the structures of the nanotubes in their primed and extended states, providing mechanistic insights into substrate exchange. Additionally, we performed comparative proteomics and genomic analyses to identify host proteomic changes in response to the DPANN attachment. Our results showed striking alterations in host-proteome during symbiosis and upregulation/downregulation of key cellular pathways. Collectively, these results provided unprecedented insights into the structural basis of host-DPANN communication and deepen our understanding of the host ectosymbiotic relationships.

## Introduction

DPANN, an acronym of the names of the first included 5 phyla (Diapherotrites, Parvarchaeota, Aenigmarchaeota, Nanoarchaeota, and Nanohaloarchaeota)^1^, is a superphylum containing more than 10 phylum-level lineages, most of which remain uncultivated. The common features of DPANN lineages are extremely small cell and genome sizes with very limited metabolic capabilities^2^. Although rare free-living DPANN lineages exists, they mostly adopt an obligate symbiotic lifestyle dependent on other microbial community members^3,4^. DPANN accounts for approximately half of all the archaeal diversity^2,5^, and are detected across the entire biosphere^3–14^ including within the human microbiome^6^. However, their biological functions, cell biology and molecular basis of symbiosis are still poorly characterised since only three representative phyla have ever been cocultured^7–13^.

One of the DPANN lineages, the phylum “*Ca.* Micrarchaeota” (or Microcaldota^14^), originally called ARMAN^15^, has been found in various natural environments (e.g., acid mine drainages (AMD), hot springs, soil, peat, hypersaline mats, and freshwater^16,17^). Since the first discovery of ARMAN in AMD^15^, its host has been generally suggested to be members of the order *Thermoplasmatales* (the phylum Euryarchaeota)^18,19^. However, we have recently established a coculture system composed of a novel Micrarchaeon “*Ca.* Microcaldus variisymbioticus ARM-1” (hereon referred as ARM-1) and its host “*Metallosphaera javensis* AS-7” (hereon referred as AS-7) belonging to the order *Sulfolobales* (the phylum Crenarchaeota)^14^. Notably, most *Thermoplasmatales* spp. lack a surface layer (S-layer)^20^, while *Sulfolobales* spp. have a defined S-layer^21^. This difference in the cell envelope, in combination with other unknown factors, could contribute to host specificity of the “*Ca.* Micrarchaeota”.

Previous studies suggested that DPANN-host interaction could involve membrane fusion and direct cytoplasmic connection (or, “cytoplasmic bridge”) between the host and the symbiont, facilitating the exchange of nutrients and even enzymes^11,18,19,22,23^. The molecular and structural basis of host-DPANN attachment, “cytoplasmic bridge” formation, and host-DPANN symbiosis remain poorly understood. Intriguingly, a few DPANNs were recently reported to engage with their hosts without the apparent presence of a cytoplasmic bridge^14,24^. How intercellular communication is facilitated in the absence of a direct cytoplasmic connection is unclear.

Recent advances in electron cryo-tomography (cryo-ET) and associated methods have enabled the investigation of cellular ultrastructures and large macromolecular machines *in situ* in their near-native state at unprecedented resolution^25–27^. Here, we used cryo-ET and complementary approaches, including proteomics and genome analysis to investigate the interaction between Micrarchaeota ARM-1 and its host AS-7 at molecular resolution. Our *in situ* structural analyses revealed a novel proteinaceous nanotube structure that bridges the cytoplasms of AS-7 and ARM-1 and might act as a conduit for molecular exchange. We resolved the structural details of the host and DPANN S-layers, and the bridging tube connecting the two cell types. Finally, our proteomic analysis revealed host response upon DPANN attachment and provided insights into the molecular basis of host-DPANN symbiosis.

## Results

### Electron cryo-tomography uncovers nanometre-resolution details of host-DPANN interaction

We utilised cryo-ET to visualize the interactions between ARM-1 and AS-7 in their near-native state, at nanometre resolution. ARM-1 was co-cultured with host AS-7 using methods previously described in Sakai *et al* ^14^, and the growth of the two cultures was monitored by qPCR using 16S specific primers (Supplementary Fig. 1A). Association between ARM-1 and AS-7 was initially assessed by Scanning Electron Microscopy (SEM) and subsequently imaged by cryo-ET (Supplementary Fig. 1B-D). Our cryo-tomograms showed multiple ARM-1 Micrarchaeota interacting tightly with each AS-7 cell (Fig. 1A,B, Supplementary Movie 1,2). The different cell types (host and DPANN) are distinguishable by the appearance of their S-layers and overall sizes; the AS-7 host cell has a characteristic corrugated S-layer and measures 1.5 to 2 µm in diameter, while ARM-1 measures 0.3 to 0.5 µm in diameter (Fig. 1A,B and Supplementary Fig. 1B-D). Density profile analysis of the ARM-1/AS-7 envelopes revealed that the distance from ARM-1 S-layer to AS-7 S-layer was between 11 and 16 nm (Supplementary Fig. 2). Close examination of AS-7 cells showed proteinaceous ‘tube-like” structures embedded just beneath the S-layer (Fig. 1C,E, yellow arrows). Intriguingly, we also observed many of these nanotubes extending between the two cell types bridging their cytoplasms (Fig. 1A,C). Nanotubes embedded within the AS-7 S-layer are likely the primed state that upon some stimulation extend to form host-DPANN contact. The number of extended nanotubes varied between different ARM-1 and AS-7 interactions with an average of 4.4 nanotubes per symbiont/host contact (Supplementary Fig. 3A,B). In some of our tomograms, we observed more than 10 nanotubes connecting each ARM-1 to its host, whereas in few others, there were none (Supplementary Fig. 3A,B).

**Figure 1.**
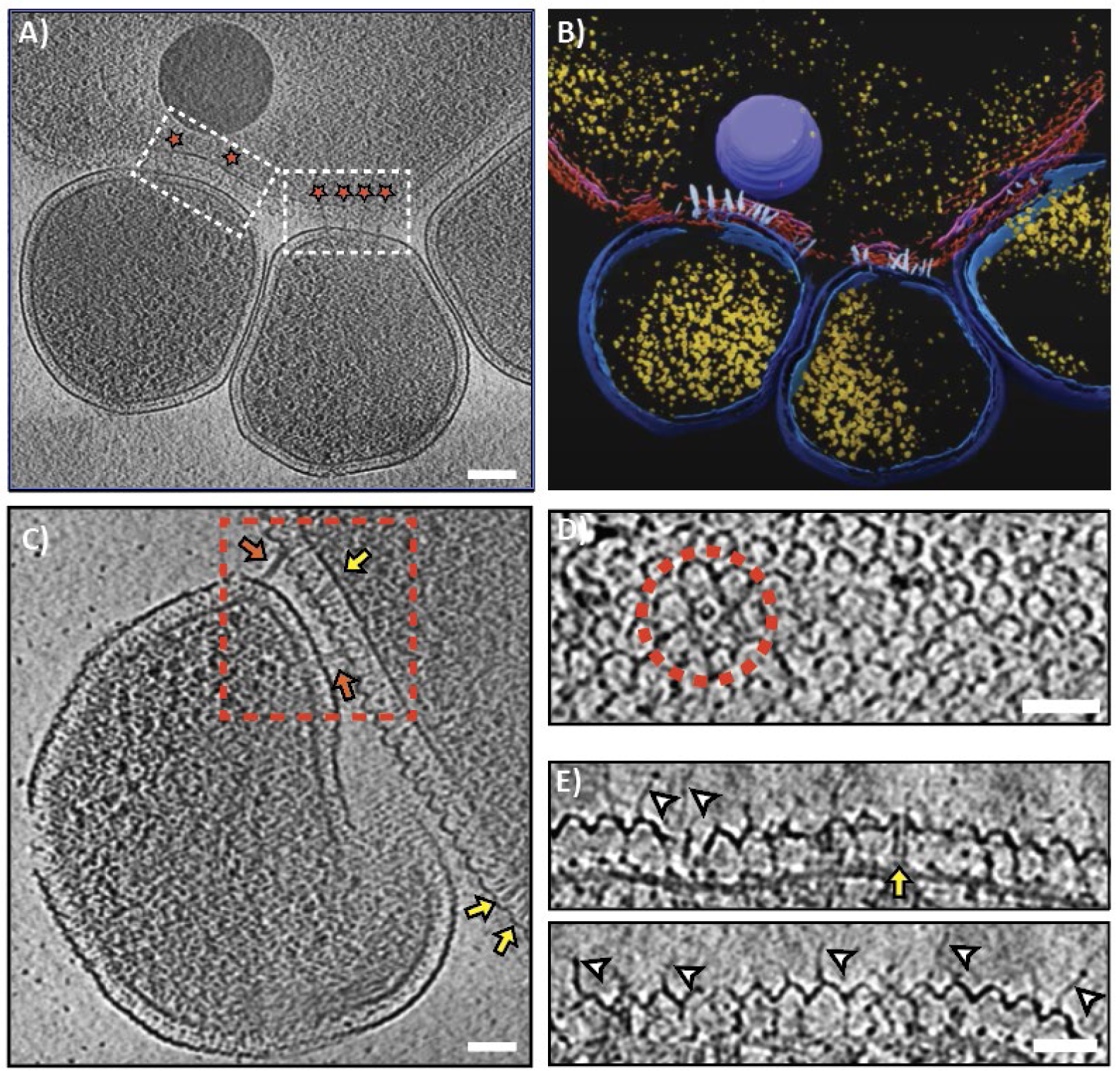
Molecular interactions of the ARM-1 – AS-7 coculture. (A) Representative tomographic slice of ARM-1 (three smaller cells) cells interacting with their host AS-7 (large single cell, top). White boxes highlight interacting interfaces of ARM-1 and AS-7, and red stars indicate the location of tube structures adjoining the two organisms. (B) Segmentation analysis of the 3D volume highlights key components: tube like structures (white), host S-layer (red), host inner membrane (pink), ribosomes (yellow), ARM-1 S-layer (dark blue), and ARM-1 cytoplasmic membrane (light blue). (C) Enlarged view of another ARM-1 cell (centered) interacting with the AS-7 host (top right). Views of tube-like structures are highlighted by a red box, inter-cell connecting tubes (red arrows) and the side view of a primed tube within the AS-7 membrane envelope (yellow arrows) are clearly visible. (D) Top view of an AS-7 cell a red dashed circle highlights the top view of a primed tube in the S-layer. (E) Two side views of the host AS-7 membrane and S-layer, thin protrusions are shown (white arrows) which are a different structure to the tubes (yellow arrow). Images in (A), (C), (D), and (E) are 2D slices through 3D reconstructed volumes generated from tilt series of 2D projection images. Scale bars indicate 100 nm (A), 50 (C-E) nm.

Top views of our reconstructed tomograms revealed that the AS-7 S-layer forms a uniform lattice with pores of P6 symmetry. The hexagonal arrangements often had a circular structure at the centre (Fig. 1D). Careful investigation of our tomograms from different axes and measurement of dimensions confirmed that these circular structures are top views of the nanotubes embedded within the AS-7 S-layer (Fig. 1D,E). Interestingly, the nanotubes were structurally aligned to the S-layer pores, suggesting that the AS-7 S-layer likely forms a guiding scaffold for the tube assembly and extension. The frequency of nanotubes varied between different tomograms and had a strikingly positive correlation with the presence of the symbiont.

In many of our tomograms, we observed several thin filament-like “antennae” densities protruding out from the S-layer of AS-7 cells into the extracellular environment (Fig. 1E, white arrows). We were unable to resolve if the antennae traversed the entire cell envelope. The tip of the antennae harboured a bulb-like density reminiscence of bacteriophage tail fibres^28^. The antennae might play a role in establishing the initial contact before host-DPANN cytoplasms are connected by nanotubes. These results show the nature of ARM-1 and AS-7 interaction in molecular detail and reveal previously uncharacterized protein nanotube and putative receptor structures that mediate host-DPANN interaction.

Intriguingly, the host-DPANN interaction showed a remarkable resemblance with the phagocytosis process. In some of our tomograms, we saw a gentle depression of host membrane near the AS-7 and ARM-1 contact site, forming a phagocytic cup-like structure (Supplementary Fig. 4A). In other tomograms, we noticed pseudopod-like extensions surrounding the ARM-1 symbiont (Supplementary Fig. 4B). Finally, we captured one interaction where an ARM-1 cell was nearly engulfed by the host (Supplementary Fig. 4C, Supplementary Movie 3). Recently, Hamm *et al* proposed a predatory, intracellular life cycle of a DPANN archaeon (*Candidatus* Nanohaloarchaeum antarcticus)^29^. Although we saw different landmarks of phagocytosis-like internalisation, we did not notice lysis of AS-7 cells on our grids, suggesting the AS-7 and ARM-1 interaction is likely ectosymbiotic and nonparasitic.

### *In situ* structural analyses of the AS-7 S-layer show protein nanotubes originate from the host

Tomographic slices parallel and perpendicular to the AS-7 S-layer revealed that the S-layer formed a 2D regular hexagonal array, which is also confirmed by the power spectrum analysis (Supplementary Fig.5A-D). To obtain a detailed understanding of the surface organisation of AS-7 and resolve the *in situ* structure of the nanotube in its primed state, we performed subtomogram averaging (STA) of the AS-7 S-layer with and without the nanotube.

For the AS-7 S-layer, we first manually picked 200 particles to produce a *de novo* model in EMAN2^30,31^. This model was then used as a reference for template matching to pick 10,711 particles from 72 tomograms. STA of the AS-7 S-layer lattice resulted in a ∼1 nm resolution structure of the S-layer, where S-layer protein (SlaA and SlaB) domains were distinctly resolved (Fig. 2A-C, Supplementary Fig. 5A,E). This allowed us to generate AlphaFold models of SlaA and SlaB and fit those in our subtomogram average map, revealing overall orientation and organization of SlaA and SlaB within the S-layer lattice (Supplementary Fig. 5F-I, Supplementary Movie 4). The local resolution of our structure was 8-16 Å as estimated by ResMap^32^ (Supplementary Fig. 6C,D). Our structure showed that the S-layer formed a regular array of dome-like structures at a centre-to-centre distance of ∼21 nm (Fig. 2C). The dome-like structure was rotationally 6-fold symmetric with a small pore in the center, which had a diameter of 3.5 nm (Fig. 2A,C).

**Figure 2.**
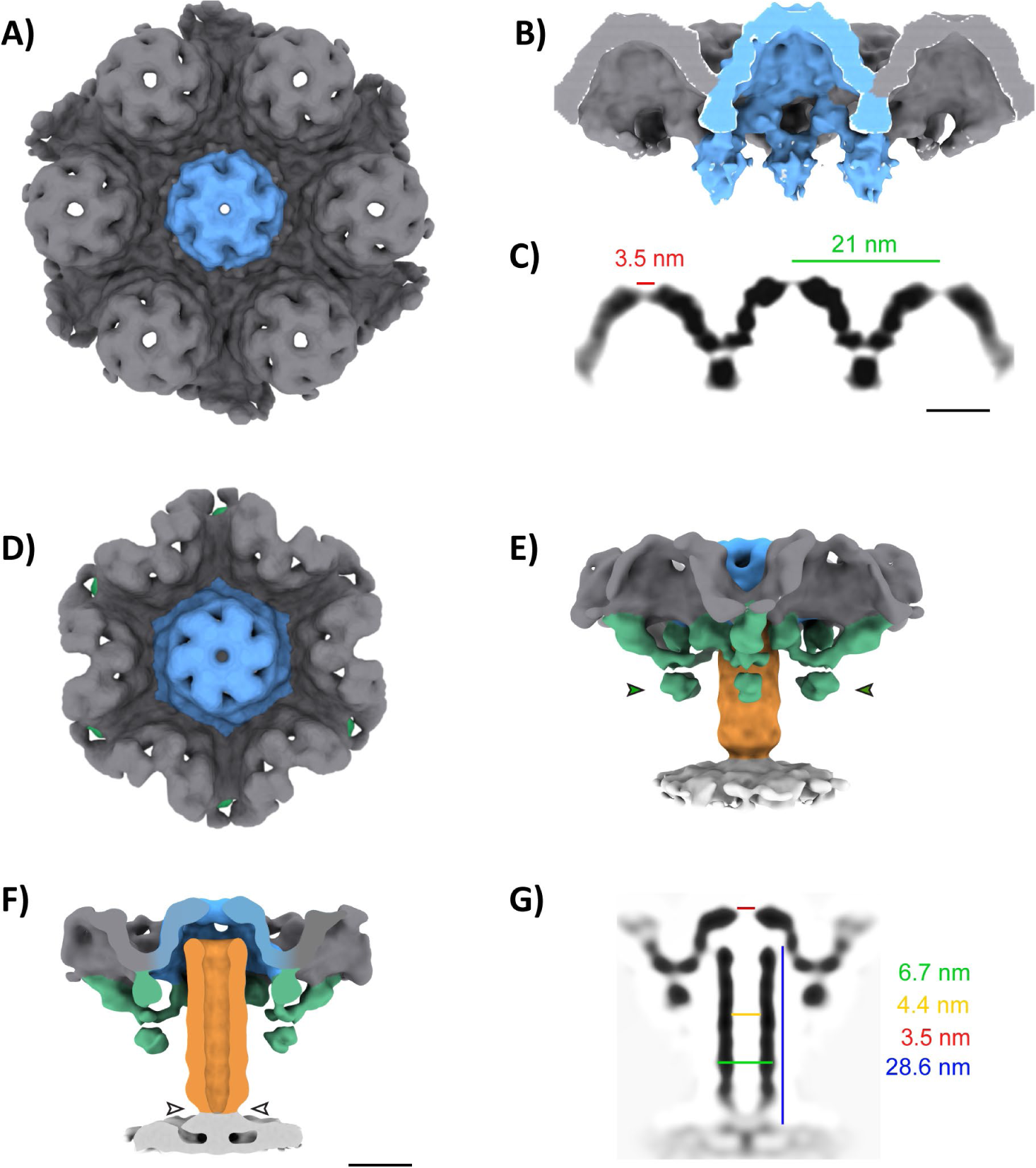
*In situ* structure of the AS-7 S-layer and primed nanotube. (A) Surface representation of the host cell AS-7 S-layer structure determined by STA (grey), an individual S-layer cap is highlighted in blue. Cross-section views of figure 2A as a surface representation (B) and orthogonal plane view (C). Surface representation of the primed tube structure determined by STA, coloured based on segmentation to show the S-layer (grey and blue), tube density (orange), putative tube accessory proteins (green) and inner membrane (white) shown from outside to inside of the cell (D) and side on (E). (F) Cross-section depiction of figure 2E. (D) 2D cross-sectional orthogonal view of figure 2E. Scale bars 20 nm.

To determine the structure of the embedded/primed protein nanotube, we manually picked 797 side and top views of nanotubes embedded within the S-layer and generated a STA (Fig. 2D-G). The local resolution of the protein nanotube structure varied between 10-20 Å, as calculated by ResMap (Supplementary Fig. 6A,B). The resulting average revealed that the envelope-spanning nanotube was assembled within the S-layer scaffold and extended from the dome to the cytoplasmic membrane (Fig. 2E-G). The nanotube was structurally aligned with the pore of the S-layer, suggesting that the S-layer pore could act as conduit for nanotube extension during host-interaction (Fig. 2G). The base of the nanotube, near the cytoplasmic membrane, was narrower suggesting there might be a gating mechanism. The top and vertical cross-section views of the average best illustrated the architecture of the membrane-spanning complex, but the overall organisation was better viewed in the segmented volume (Fig. 2E-G, Supplementary Movie 5). The segmented volume highlighted 6 necklace-with-pendant-like densities around the tube, extending down from the S-layer (Fig. 2E), providing a further framework for the nanotube structure. The length of the nanotube from the dome to the cytoplasmic membrane was 28.6 nm and the inner and outer diameters were 4.4 and 6.7 nm respectively, suggesting a wide-enough lumen for metabolite and unfolded protein transport (Fig. 2G). Given the AS-7 S-layer pore diameter is 3.5 nm, the nanotube extension cannot occur without a large conformational change within the S-layer or a symmetry break. Since nanotubes were also present in AS-7 pure culture (Supplementary Movie 6) and were integrated within the AS-7 S-layer, the connecting tubes originate in the AS-7 host cells and not the ARM-1 DPANN.

### *In situ* structure of the AS-7 extended nanotube

To understand the structural basis of host-DPANN interaction, we attempted to generate an STA of the extended tube connecting cells. Our initial attempts to average the extended tube using EMAN2 were unsuccessful due to the considerable range of angles that the tube protruded from the AS-7 envelope (Supplementary Fig. 7). Therefore, we adopted a guided approach where relative orientations for each particle within cryo-tomograms were provided as *a priory* information for further focused alignment and refinement. In our initial average, part of the tube entering the ARM-1 envelope resolved well but densities around the AS-7 S-layer were lacking. Therefore, we used masks to perform focused alignment and averaging of the tube near the ARM-1 envelope separately from that of the AS-7 envelope. By combing the two averages, we generated a composite STA of the intercellular nanotube bridging the DPANN and host (Fig. 3A,B). Our structure showed that the nanotube originates in the AS-7 cytoplasmic membrane and traverses through the AS-7 and ARM-1 S-layers, connecting their cytoplasms (Fig. 3A,B). The extended nanotubes are 68 nm long, with an outer and lumen diameter of 6.7 and 4.4 nm, respectively (Fig. 3B). Intriguingly, we observed a barrel-like structure in the ARM-1 envelope seemingly acting as a docking platform for the in-coming tube. The barrel-like structure is reminiscent of the type-II secretion system secretin complex but with a much larger diameter of 21 nm (Fig. 3B)^33^. Because the tubes emanated at diverse angles from the AS-7 envelope, despite a guided approach and focused alignment, the AS-7 S-layer resolved poorly in our composite average. (Fig. 3B, Supplementary Fig. 7).

**Figure 3.**
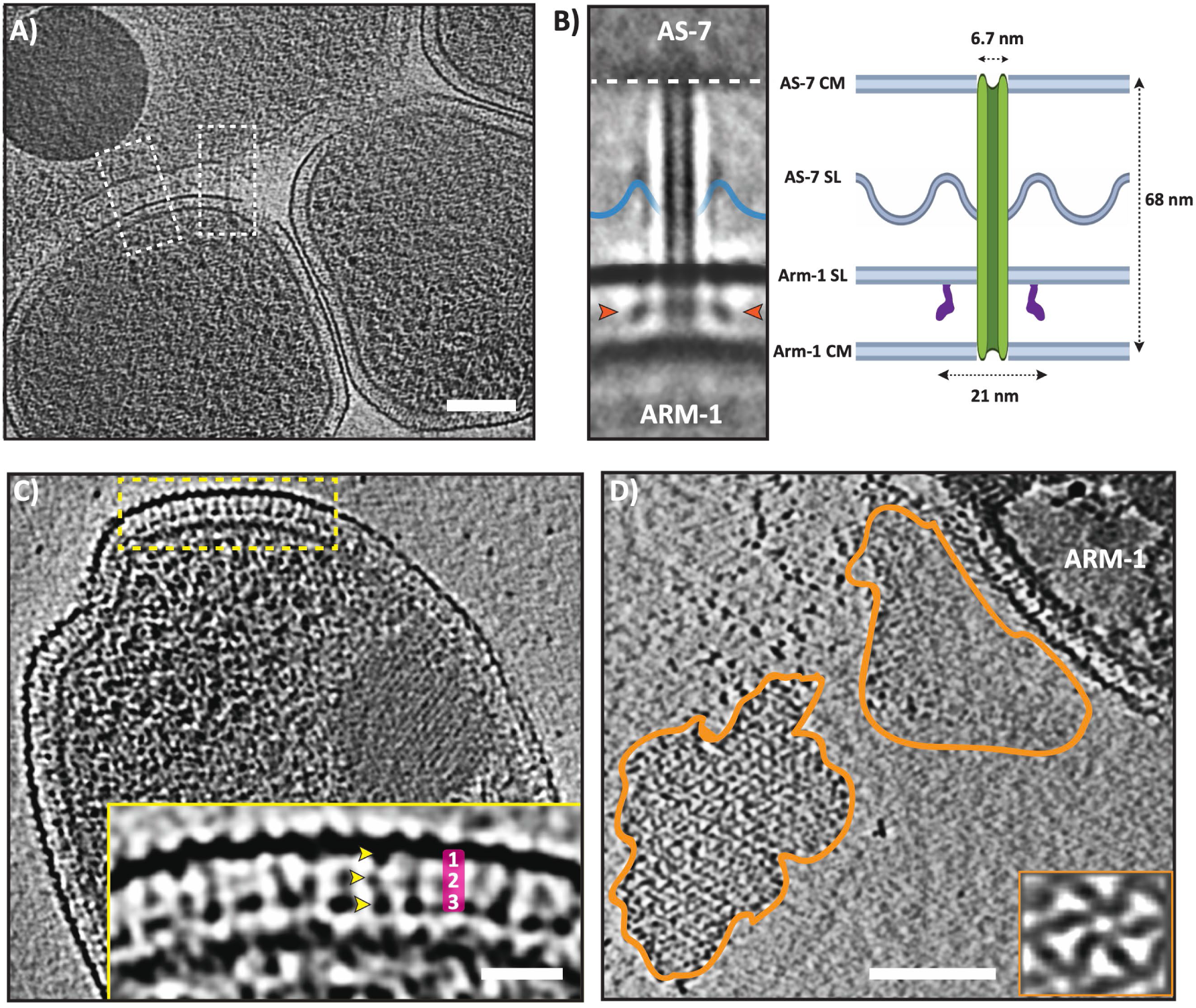
Architecture of the DPANN (*C. Micrarchaeota)* S-layer and extended intercellular nanotube. (A) Tomographic slice of a DPANN-host assembly showing intercellular tubes (white boxes) between DPANN and its host. (B) A composite subtomogram average of the intercellular tube between DPANN and its host. The intercellular tube originates in AS-7 and extends into DPANN. Beneath the DPANN S-layer, a large barrel-shaped complex (red arrowheads) forms a platform for the incoming nanotube. The position of the AS-7 S-layer is depicted based on the weak density visible on the average. A schematic of the intercellular tube with distances/diameters shown on the right. (C) Tomographic slice of a DPANN cell. Side-view of the S-layer showing distinct structural features (yellow box). Inset showing magnified side-view of the S-layer. Crescent-shaped protein densities are seen extending from the outer S-layer to the cytoplasmic membrane. At least 3 globular domains are visible (yellow arrowheads, numbered). nm. (D) Tomographic slice of a DPANN cell with partially detached/peeled off S-layer. Subtomogram averaging of the top-views showed 6-fold symmetry. Scale bar (A, D) 100 nm, (C) 25 nm.

### The ARM-1 outer membrane is comprised of an S-layer

No structural elucidation of the S-layer has been achieved from any of the DPANN superphylum members to date. Our attempts to generate an STA of the ARM-1 S-layer using multiple different software were unsuccessful. This could be because of the highly bent nature of the ARM-1 S-layer (due to small size and curvature) and/or imperfect lattice arrangement. Unlike the host AS-7 S-layer, no corrugated dome-like arrangement was observed in the ARM-1 envelope and only one ortholog for SlaA was found in the ARM-1 genome, suggesting a different S-layer organization. Side views of the ARM-1 envelope showed crescent-shaped structures, extending downwards, with three discrete densities extending from the outermost layer towards the cytoplasmic membrane (Fig. 3C, inset). The distribution and orientation of these densities were visibly irregular, suggesting imperfect lattice arrangement. In a few tomograms of partially lysed ARM-1 cells, S-layers were found partly detached from the main cell body (Fig. 3D). Recent studies have used methods to artificially shed S-layers from cells using detergents in order to obtain native structures of S-layers^34^. We used this opportunity to generate an average of the ARM-1 S-layer top view (Fig. 3D, inset). Our average showed that the ARM-1 S-layer constitutes an ordered 2D S-layer lattice of 6-fold symmetry with a pore in the centre. The centre-to-centre distance between neighbouring hexametric arrangements was 18.5 nm.

### Comparative proteomics of the host and coculture reveals molecular players involved in the **symbiotic relationship.**

Our cryo-ET analysis revealed that the host archaeal cell AS-7 interacts with the ARM-1 symbiont through intercellular nanotubes, and that these tubes originate and extend from the AS-7 host. To gain insights into host response to ARM-1 interaction and identify plausible resources being exchanged between the host and the symbiont, we performed comparative proteomics of pure host and host-DPANN cocultures. Using Label free quantification (LFQ) based proteomics a total of 698 ARM-1 proteins and 1810 AS-7 proteins were identified with the presence of ARM-1 resulting in widespread alterations within the AS-7 proteome (Fig. 4A-B). Of the 698 proteins identified from ARM-1, 637 were robustly identified across all samples with sequence analysis demonstrating much of the observed proteome lacks identifiable PFAM domains or similarities to known proteins based on GO and KEGG assignments (Supplementary Fig. 8A). Despite this, analysis of the relative abundance of the observable proteome based on iBAQ analysis^35^ (Supplementary Fig. 8B) supports that the observable proteome spans nearly 5 orders of magnitude and reveals proteins associated with ATP synthase are highly abundant, consistent with the active growth of ARM-1. Within the observable AS-7 proteome, 134 proteins were significantly increased in abundance while 373 were decreased upon ARM1 interaction (Fig. 4A-C). Overlaying these observed proteomic changes on the genome revealed multiple loci to display congruent changes within neighbouring gene products (Fig. 4 D-F, Supplementary Table 1). For example, the addition of ARM1 resulted in an increase in multiple gene products of BCS91585…1-BCS91596.1 and BCS93590.1…BCS93624.1, with these proteins associated with CO_2_ fixation, and the TCA cycle; BCS92975.1…BCS92983.1, encodes proteins that are predicted to be involved in inorganic ion transport and metabolism (Fig. 4E, cluster i); and BCS93537.1…BCS93550.1 involved in aromatic metabolism, tyrosine metabolism and branched chain amino acid metabolism (Fig. 4F, cluster ii). Comparative genomics analysis of these impacted AS-7 proteins supports that cluster (ii) is found exclusively in AS-7 suggesting a role for this locus for the AS-7 and ARM-1 interaction (Supplementary Fig. 9, Supplementary Table 2).

**Figure 4.**
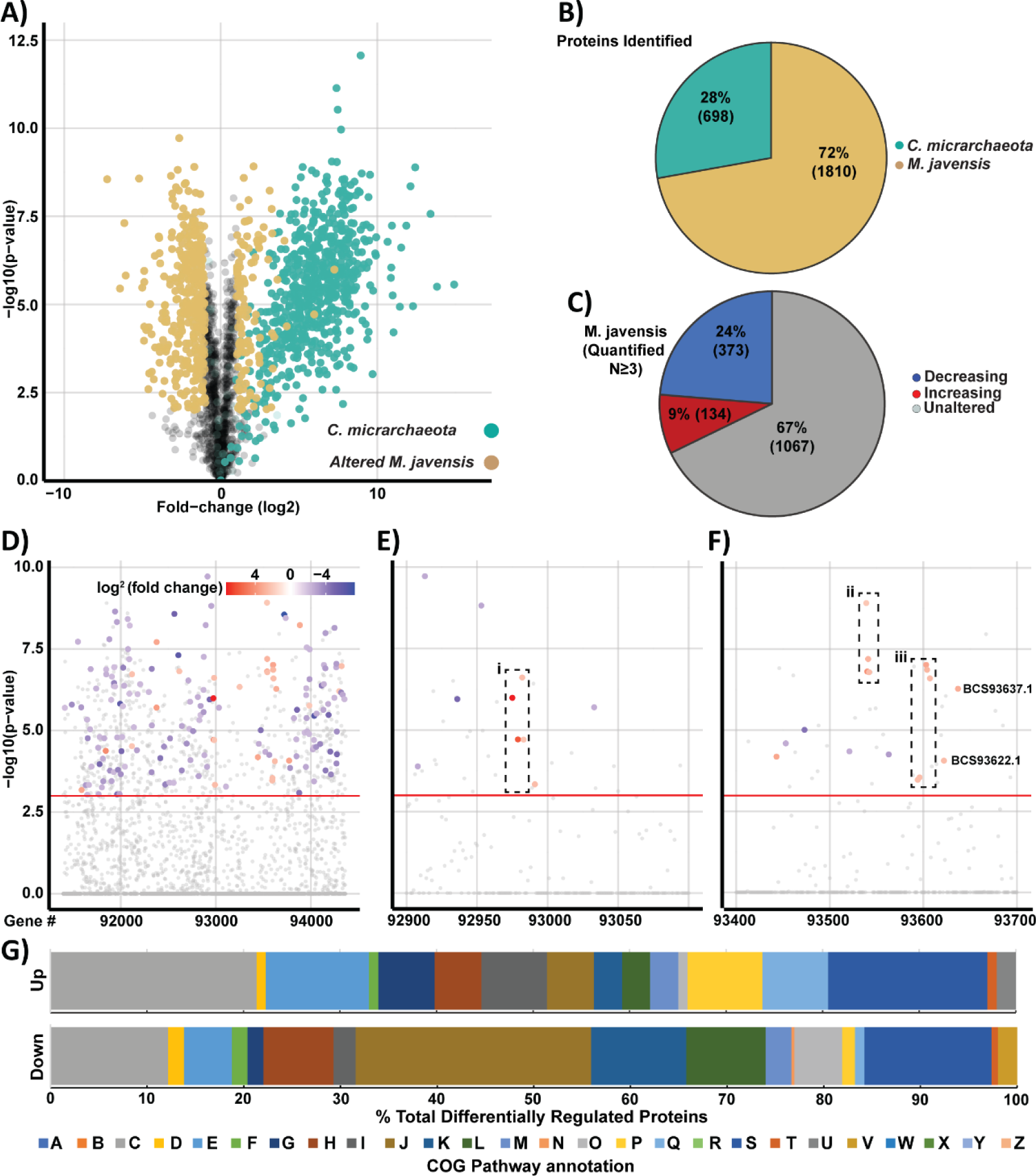
Proteome analysis of *C. Micrarchaeota* and *M. javensis*. A) Volcano Plot of proteome alterations observed between *M. javensis* co-cultured with *C. Micrarchaeota* and *M. javensis* alone. Proteins observed to be altered within *M. javensis*, defined as proteins showing a > ±1 log2 fold change and a -log10(p-value)> 2, are denoted in yellow while *C. Micrarchaeota* proteins are denoted in green. *C. Micrarchaeota* proteins are presented here to show their identification, not fold change. B) Pie chart of observable proteins across species reveal 698 and 1810 proteins were identified across these proteomes respectively at a 1% FDR. C) Pie chart of proteins quantified ≥3 replicates of *M. javensis* (1654 proteins) reveal 33% of the proteome is altered during *C. Micrarchaeota* infection. D) Manhattan plot of protein changes observed across the *M. javensis* genome reveals widespread alterations with examination of protein changes between accession BCS92900 and BCS93100 (E) and BCS 93400 and BCS93700 (F) revealing evidence of alterations within potential operons. Bar graph of COG pathway analysis of proteomics data, showing the percentage of differentially regulated proteins in each COG category (G). Standard COG categories (A-Z) are as follows: RNA processing and modification (A), Chromatin Structure and Dynamics (B), Energy production and conversion (C), Cell cycle control, cell division, chromosome partitioning (D), Amino acid transport and metabolism (E), Nucleotide transport and metabolism (F), Carbohydrate transport and metabolism (G), Coenzyme transport and metabolism (H), lipid transport and metabolism (I), Translation, ribosomal structure and biogenesis (J), Transcription (K), Replication, recombination and repair (L), Cell wall/membrane/envelope biogenesis (M), Cell motility (N), Posttranslational modification (O), Inorganic ion transport and metabolism (P), Secondary metabolites biosynthesis, transport and catabolism (Q), General function prediction only (R), Function unknown (S), Signal transduction mechanisms (T), Intracellular trafficking, secretion, and vesicular transport (U), Defense mechanisms (V), Extracellular structures (W), Mobilome: Prophages, transposons (X), Nuclear structure (Y), and Cytoskeleton (Z)

COG analysis of the 373 significantly decreased proteins in the AS-7 reveals that most differentially down regulated proteins appear involved in translation, ribosomal structure and biogenesis (COG:J), transcription (COG:K), and replication, recombination and repair (COG:L) (Fig. 4 G). Specifically, a four gene operon (BCS92057.. BCS92060) encoding proteins for the exosome complex, which is responsible for RNA degradation; a four gene operon (BCS91961..BCS91964), containing three hypothetical proteins that have predicted structures similar to SNARE proteins that are involved in vesicle formation and trafficking; a six gene operon (BCS94267..BCS94277) containing Alba DNA binding proteins and DNA topoisomerases. Collectively, these results indicated that the AS-7 cell is reducing its proliferation rate in response to ARM-1 interaction.

### Filamentous structures, protein-cages and large membrane-tubes between ARM-1 cells

Interestingly, in some of our tomograms, ARM-1 cells had stacks of filament-like structures close to the membrane, resembling filamentous phages^36^ (Supplementary Fig. 10A). We also noticed, large protein-cage-like structures that were often more than 100 nm long and 50-70 nm wide assuming barrel or trapezoid-like shapes (Supplementary Fig. 10B, Supplementary Movie 2). These assemblies are similar to those previously reported^37^, despite the improved resolution presented here, their function remains unknown. In addition, we observed thin filaments protruding out of ARM-cells that could represent archaeal pili (Supplementary Fig. 10C). Finally, in a few of our tomograms, we saw two ARM-1 cells were connected by wide membrane tubes (Supplementary Fig. 10D-F, Supplementary Movie 7). Such nanotubes/nanopods have been previously described in Crenarchaeota and Euryarchaeota but their exact function remain unclear^38^.

## Discussion

Here, using complementary approaches, including *in situ* structural biology, proteomics and genomic analysis, we investigated the structural and molecular basis of the episymbiotic relationship between *Candidatus Micrarchaeota* ARM-1, a DPANN archaea, and its archaeal host *M. javensis* AS-7. Our results characterize the host-symbiont interactions at molecular resolution revealing an antenna-like structure on the surface of the host, and an intercellular nanotube structure, bridging the host and DPANN cytoplasms. The antenna structure could mediate symbiont recognition and initial contact formation, leading to the biogenesis of the nanotube apparatus. Previous low-resolution studies have shown that DPANNs can form different types of interactions with their hosts including: cytoplasmic bridges^18^, parasite-like invasion^11,39^ , and pili-like connections^18,40^. Here, we discovered a novel nanotube structure that bridges the host and symbiont, promoting intercellular exchange. Our nanotube structures have a resemblance to previously described connections between *Thermoplasmatal* and associated DPANN (ARMAN). Due to limited resolution, these connections were referred to as “filamentous connections”^18,40^. *Thermoplasmatal* archaea are distantly related to *Metallosphaera* indicating that these nanotubes maybe conserved across different archaea. Our cryoelectron tomograms revealed that the nanotubes are in two different conformational states. Firstly, a primed state where the nanotube is assembled and embedded within the AS-7 cell envelope, beneath the S-layer. Second, an extended state where the nanotube elongates and traverses through the symbiont envelope and establishes cytoplasmic connection between the two cell types. The primed tubes were also present in host only culture, but at much lower frequency. This suggested that the nanotube originates in the host and its biogenesis is upregulated in the presence of the symbiont. A composite average of the extended tubes confirmed that the extended nanotubes penetrate into the ARM-1 cytoplasm, supporting the notion that these tubes represent a mechanism for exchange of cytoplasmic components between the host and symbiont. Although primed nanotubes are structurally aligned to the S-layer pores, the diameter of the tubes are larger compared to S-layer pore size, suggesting large-scale structural rearrangement and/or symmetry breakage in the S-layer pore is essential for the tube extension. Intriguingly, we discovered a large barrel-shaped structure in the ARM-1 cell envelope that acts as a docking platform for the incoming nanotube. The barrel-shaped structure is reminiscent of bacterial secretin-like assemblies but much larger in diameter (21 nm). The structure could be formed from AS-7 proteins secreted through the tubes that assemble in the ARM-1 envelope. Considering the symbiotic nature of the interaction, an alternative hypothesis is that ARM-1 proteins could coordinate with the extended tube to build this docking platform. The protein components that comprise the main tube structure, and any accessory proteins are the subjects of ongoing research.

Employing cryo-ET and STA, we were able to determine a 1 nm resolution *in situ* structure of the host S-layer, where domains of S-layers proteins were distinctly visible. The *in situ* structure of the AS-7 S-layer revealed a similar architecture to previously characterized S-layers from *Sulfolobus islandicus*, a related Archaea of the same family^41^. The AS-7 host S-layer constitutes an array of dome-like structures with a pore in the centre. Nanotube extension could happen through this pore but would require conformational rearrangement as the tubes are larger in diameter. Different conformations of SlaA have been reported in *Sulfolobus spp* that alter the S-layer lattice in response to the environment^41^. However, conformational changes that enlarge the pore are yet to be demonstrated *in situ*.

Interestingly, the AS-7 and ARM-1 interaction exhibited remarkable similarity and membrane dynamics to the process of phagocytosis of intracellular pathogens. In many of our tomograms, we found the AS-7 envelope displaying a phagocytic cup-like structure at the ARM-1 contact site, pseudopod-like structures extending from the host around the ARM-1 symbiont, and finally, there was one tomogram (out of ∼90) where an ARM-1 cell was nearly engulfed by an AS-7 cell. Recently, Hamm *et al* proposed an intracellular parasitic (aka, predatory) lifestyle of the DPANN archaeon *Candidatus* Nanohaloarchaeum antarcticus. It is possible that ARM-1 also has an intracellular life-cycle. However, this would be rare and nonparasitic, as we did not see lysis or internalisation of AS-7 cells on our grids.

DPANNs are characterized by their small size and small genomes (0.5 to 1.5 Mb), which often lack genes for essential metabolic processes such as amino acid and nucleotide synthesis^42–45^. The genome of ARM-1 was previously shown to be deficient in genes for glycolysis, gluconeogenesis, carbon and nitrogen fixation, and the biosynthesis of amino acids and nucleotides^14^. The reliance of ARM-1 and other DPANNs on their hosts is therefore evident, whereas the benefit to the host cell in this symbiotic relationship is less obvious. The ARM-1 AS-7 co-culture system provides an example of a host committing substantial resources and adaptations to facilitate an interaction with a DPANN, which places new emphasis on the importance of this interaction for the host cell.

STA of the nanotube showed an inner lumen diameter of 4.4 nm which is sufficiently wide to allow for folded larger globular proteins (100-150 kDa), DNA, and metabolites to pass ^46,47^. DPANN have been suggested to exchange nutrients, including amino acids, nucleotides, and sugars (for cell wall synthesis and metabolism). Confirmation of the molecular identity of the nanotube apparatus and substrates passing through the tube is difficult in our genetically intractable systems. Therefore, we utilised proteomics approaches to investigate molecular players involved in the AS-7 and ARM-1 symbiosis. Our proteomics analysis indicated that CO_2_, TCA cycle, and amino acid metabolic pathways were upregulated in the host, these data suggest that metabolites from these processes are the intended cargos. In addition, CRISPR Cas9 proteins were upregulated in the host and DPANN during co-culture, a locus containing three hypothetical proteins were shown to be an operon associated with a CRISPR Cas9 system (BCS94299…BCS942301) (Supplementary Table 1). A recent study has uncovered the important role that CRISPR systems have in host-DPANN symbiosis; it is therefore plausible that the tubes may facilitate the transfer of DNA^48^.

Cryo-ET can provide high resolution cell biology insight into an otherwise inexplorable group of organisms. We surveyed the ARM-1 cells for cellular structures that may prove important for understanding their biology and the nature of their symbiosis, our results are presented as a gallery of features that we believe will be of importance to other researchers in the DPANN field (Supplementary Fig. 10). In addition, membrane tubes (as opposed to the protein tubes discussed above) connecting DPANN cells were observed in some tomograms that are suggestive of intercellular nanotubes. Intercellular nanotubes/nanopods have been shown in archaeal cells before, however their function remain enigmatic. DPANN archaea are co-located on the host cell creating a micro community. Nanotube formation is a possible method to exchange nutrients within the DPANN community.

The world of DPANNs remain largely unexplored, their small genomes and obligate symbiotic lifestyles make their biology a fascinating area of research. The nature and evolution of these relationships is complex, and as we have shown here, the host can spend substantial resources to establish interaction and exchange biological molecules with the DPANN. At least in our system host-DPANN relationship appears to be mutualistic and ectosymbiotic. Our work demonstrates the capability of cryo-ET and STA to interrogate these genetically intractable systems and contributes to the expanding knowledge of DPANN biology and their symbiotic relationships.

## Acknowledgements

This project was funded by an NHMRC grant (APP1196924 to DG), an HFSP grant (RGEC33/2023) to DG and HDS, The Japan Society for the Promotion of Science through Grants-in-Aid for Scientific Research (21K15153 to HDS). DCS is supported by the Melbourne Research Scholarship. We are thankful to Zhenan Li for help with data processing. N.E.S is supported by an Australian Research Council Future Fellowship (FT200100270) and an ARC Discovery Project Grant (DP210100362). Cryo-EM data was collected at the Ian Holmes Imaging Centre, Bio21, University of Melbourne. We thank the Melbourne Mass Spectrometry and Proteomics Facility of The Bio21 Molecular Science and Biotechnology Institute for access to MS instrumentation.

## Data availability

Subtomogram averages of the AS-7 S-layer and primed nanotube are available under accession codes EMD-42578 and EMD-42579. The mass spectrometry proteomics data have been deposited to the ProteomeXchange Consortium with the dataset identifier PXD045678.

## Competing interests

The authors have no competing financial interests.

## Materials and Methods

### Cultivation conditions and growth monitoring

A coculture system composed of *Microcaldus variisymbioticus* strain ARM-1 and *Metallosphaera javensis* strain AS-7 was established as described previously ^14^. The coculture system, as well as pure culture of *M. javensis* were cultivated at 60°C under aerobic conditions. The growth of both strains was monitored by qPCR following the method as described previously ^14^.

### Cultivation conditions for cryoTEM

Cultures of *Metallosphaera javensis* AS-7 and a co-culture system composed of *Microcaldus variisymbioticus* ARM-1 and its host *Metallosphaera javensis* AS-7 were routinely cultivated at 55°C using modified Brock’s basal salt medium (Kurosawa et al. 1998) supplemented with 1 g/L yeast extract (MBSY medium) as described previously (Sakai et al. 2022. PNAS). The pH of the medium was adjusted to 3.0 by the addition of 50 % (v/v) sulfate. When the growth entered the stationary phase, the co-culture was filtered by a sterilized syringe filter (pore size: 0.45 μm, polyethersulfone). A 2 mL aliquot of the filtrate containing approximately 10^7^ cells of ARM-1 was added to 8 mL of a MBSY medium, followed by inoculation with 50 μL of pure AS-7 culture. This co-culture system was incubated at 60°C under aerobic conditions in triplicate. After 5 days of incubation, the co-culture was used for cryo-EM, 200 μL aliquots of the cultures were regularly sampled for growth monitoring by quantitative PCR analysis.

### Growth monitoring by qPCR

To monitor the growth of ARM-1 and AS-7, qPCR was conducted following the same method as described previously (Sakai et al. 2022. PNAS). Briefly, microbial DNAs of the cultures were extracted by a DNA extraction machine (magLEAD 6gC; Precision System Science) with DNA extraction reagent (MagDEA Dx SV; Precision System Science), qPCR was done with specific primers targeting the 16S rRNA gene of each species.

### Cultivation conditions for proteomics and Scanning electron microscopy

A co-culture prepared by the same procedures described above and pure AS-7 culture were pre-incubated at 55°C for 10 days. A 1 mL aliquot of the co-culture or pure AS-7 culture was inoculated into 100 mL of MBSY medium and incubated at 60°C under aerobic conditions. A total of 6 sets of 100 mL scale cultures were prepared. After five days of incubation, these cultures (100 mL x 6) were used for proteomics, whereas 5 mL from one of the co-cultures was sampled and used for scanning electron microscopy.

### Scanning electron microscopy

Microbial cells of the co-culture were fixed by 1% glutaraldehyde for 15 min at room temperature. A 5 mL of the fixed co-culture was filtered on a polycarbonate membrane filter with 0.1 μm pores (VCTP, 25mm, Merck Millipore). Cells trapped on the filter were washed by passing through 500 μL of pure water five times. After washing, the filter was flash-frozen, dehydrated, and sputter coated with osmium by the method described as previously^14^. The sputter-coated sample was analyzed at an accelerating voltage of 4 kV under the scanning electron microscope (JSM-7500F; JEOL).

### Sample preparation for Proteomic analysis

Cell suspensions were adjusted to 4% SDS, 100 mM Tris pH 8.5 and boiled for 10 min at 95 °C. The protein concentrations were then assessed by bicinchoninic acid protein assays (Thermo Fisher Scientific) and 200 μg of each biological replicate prepared for digestion using Micro S-traps (Protifi, USA) according to the manufacturer’s instructions. Briefly, samples were reduced with 10mM Dithiothreitol for 10 mins at 95°C and then alkylated with 40 mM iodoacetamide in the dark for 1 hour. Samples were acidified to 1.2% phosphoric acid and diluted with seven volumes of S-trap wash buffer (90% methanol, 100 mM Tetraethylammonium bromide pH 7.1) before being loaded onto S-traps and washed 3 times with S-trap wash buffer. Samples were then digested with 2μg of Trypsin (1:100 protease:protein) in Tetraethylammonium bromide pH 8.5 overnight at 37°C before being collected by centrifugation with washes of 100 mM Tetraethylammonium bromide, followed by 0.2% formic acid followed by 0.2% formic acid / 50% acetonitrile. Samples were dried down and further cleaned up using C18 Stage 1,2 tips to ensure the removal of any particulate matter^49,50^.

### Reverse phase Liquid chromatography–mass spectrometry

C18 enriched proteome samples were re-suspended in Buffer A* (2% acetonitrile, 0.01% trifluoroacetic acid) and separated using a two-column chromatography setup composed of a PepMap100 C18 20-mm by 75-µm trap (Thermo Fisher Scientific) and a PepMap C18 500-mm by 75-µm analytical column (Thermo Fisher Scientific) using a Dionex Ultimate 3000 UPLC (Thermo Fisher Scientific). Samples were concentrated onto the trap column at 5 µl/min for 6 min with Buffer A (0.1% formic acid, 2% DMSO) and then infused into an Orbitrap 480™ (Thermo Fisher Scientific) at 300 nl/minute via the analytical column. Peptides were separated by altering the buffer composition from 3% Buffer B (0.1% formic acid, 77.9% acetonitrile, 2% DMSO) to 23% B over 59 minutes, then from 23% B to 40% B over 10 minutes and then from 40% B to 80% B over 5 minutes. The composition was held at 80% B for 5 minutes before being returned to 3% B for 10 minutes. The Orbitrap 480™ Mass Spectrometer was operated in a data-dependent mode automatically switching between the acquisition of a single Orbitrap MS scan (300-1600 m/z, maximal injection time of 25 ms, an Automated Gain Control (AGC) set to a maximum of 300% and a resolution of 120k) and 3 seconds of Orbitrap MS/MS HCD scans of precursors (normalise collision energy of 30%, a maximal injection time of 22 ms, a AGC of 75% and a resolution of 15k).

### Proteomic data analysis

Identification and LFQ analysis were accomplished using MaxQuant (v2.0.2.0)^51^ using the *M. javensis* MjAS7 (NCBI GCA_021654415.1) and the C. Micrarchaeota ARM-1 (NCBI GCA_021654395.1) proteomes with Carbamidomethyl (C) allowed as a fixed modification and Acetyl (Protein N-term) as well as Oxidation (M) allowed as variable modifications with the LFQ and “Match Between Run” options enabled. The resulting data files were processed using Perseus (version 1.6.0.7)^52^ with missing values imputed based on the total observed protein intensities with a range of 0.3 σ and a downshift of 1.8 σ and statistical analysis undertaken using a two-tailed unpaired T-test. The mass spectrometry proteomics data have been deposited to the ProteomeXchange Consortium via the PRIDE^53^ partner repository with the dataset identifier PXD045678

### Gene conservation analysis

Conservation analysis of up-and down-regulated proteins was performed by tBLASTn of protein sequences against a curated set of Archaeal genomes from family Sulfolobaceae from the genome taxonomy database (gtdb.ecogenomic.org). These genomes were classified and checked for quality in a previous study^54^. and a full list is provided in supplementary table 2. Putative orthologues retrieved by tBLASTn were retained if they had >=50% amino acid identify and >=75% coverage relative to the query, and the presence/absence matrix generated in R. Extended annotation of the host and symbiont protein-coding sequences (from genbank accessions AP024486.1 and AP024487.1) to assign COG categories and predicted pathways was completed with EGGNog mapper version 2.1.9 with no taxonomic restriction on orthologs, and the following thresholds: evalue 0.001, score 60, % ID 40, query and subject coverage 40%.

### Electron cryotomography Data collection

Co-cultures were mixed with 10 nm colloidal gold beads (Sigma-Aldrich, Australia), which had been precoated with 1% BSA. The co-culture-colloidal gold bead mixture was added to glow-discharged copper R2/2 Quantifoil holey carbon grids (Quantifoil Micro Tools GmbH, Jena, Germany). Grids were blotted (under 100% humidity conditions) and plunge-frozen in a liquid ethane mixture (Vitrobot Mark IV, FEI Thermo Fisher Scientific). Grids were imaged using an FEI Titan Krios G4, 300 keV FEG transmission electron microscope (Thermo Fisher Scientific), micrographs were collected using a Gatan K3 Summit direct electron detector equipped with an energy filter set to 20 eV. Tilt series of cells were collected automatically between −60° to +60° at 2° intervals using the FEI Tomography 5 data collection software. The cumulative total dosage used was 100 e^−^ Å^−2^, with a defocus of −6 μm and a pixel size of 3.4 Å.

### Data processing, segmentation and sub-tomogram averaging

Tilt series were aligned with either IMOD etomo-tomo3D-PEET (Data used in segmentation, and extended tube STA) or EMAN2 (contracted tube STA, S-layer STA for host and DPANN). IMOD processing alignment used 2K binned micrographs, which were reconstructed by Tomo3D into 600 nm thick tomograms in 2K. The reconstructions were further binned to 1K for particle picking and segmentation. Segmentations were performed with the help of https://ariadne.ai/. Briefly, deep convolutional neural networks (CNNs) were trained using the manually generated ground truth and refined until a good quality of segmentation was reached. Subsequently, automated segmentations in 3D were performed. The segmentations were proofread and polished by expert inspection using the open-source software Knossos (https://knossos.app/). For each segmentation class, meshes and binary Tiff masks were generated for subsequent rendering and visualization in Knossos and Blender (https://www.blender.org/). Picked particles were then extracted from the 2K reconstructed and averaged using PEET.

In EMAN2 an initial model of the AS-7 S-layer was generated by averaging 200 high-quality S-layer particles manually selected from five tomograms. The selected particles were extracted with a box size of 52 from the tilt series at x4 binning (13.56 Å/pix). The extracted particles were then used to create a de novo initial model of the AS-7 S-layer using the initial model generator program in EMAN2.

The de novo initial model of the AS-7 S-layer was used as a reference for template matching which yielded 10,711 particles from 72 tomograms. The particles selected by template matching were extracted from each tilt series with a box size of 52 at x4 binning (13.56 Å/pix). The sub-tomograms were contrast transfer function (CTF) corrected at the per-particle-per-tilt level during extraction. The sub-tomograms were then aligned and averaged, to obtain initial alignments which were then used to re-extract particles from each tilt series with a box size of 208 without binning (3.39 Å/pix).

For particles corresponding to the primed state of the tube, located beneath the AS-7 S-layer, 797 particles were manually selected from 28 tomograms. The selected particles were extracted with a box size of 168 (3.39 Å/pix) and CTF corrected. A *de novo* model of the tube was obtained using 50 high-quality particles using the EMAN2 initial model generator.

For both S-layer and tubes (primed state) averaging, seven iterations of sub-tomogram refinement were used, consisting of three rounds of 3D particle orientation, two rounds of 2D subtilt translation, one round of subtilt translation & rotation, and one round of subtilt defocus refinement with either c1 or c6 symmetry applied. The threshold was set to exclude the worst 20% of particles at each iteration step.

### Modelling fitting and refinement

The full-length sequence of S-layer protein A (SlaA) and B (SlaB) of *Metallosphaera javensis* was predicted using AlphaFold v2.0^55^. AlphaFold multimer was used to predict the homotrimer of SlaB. The predicted models were fitted and refined into the density map representing both SlaA and SlaB proteins. Flexible N-terminal residues of SlaA (1-27) and SlaB (1-40) were removed during density fitting. Following visual inspection, the predicted full-length model of SlaA was split into 4 units (U1: 28-373; U2: 374-506, 507-639; U3: 640-784, 785-919; U4: 920-1076, 1077-1230, 1231-1345) each containing one or more domains. A semi-automated approach was employed to fit and refine individually units in a stepwise manner into the ∼10 Å resolution map of S-layer complex. Initially, the individual units were fitted and refined as rigid bodies using ‘matchpt’ and ‘collage’ tool respectively from the Situs package^56,57^. Fits were finally refined using RIBFIND and Flex-EM conjugate gradient minimisation^58,59^. For SlaB, using visual inspection, the bounding box corresponding to a single SlaB trimer was identified and extracted from the full map of the S-layer. Situs ‘matchpt’ program was used to fit SlaB trimer. For SlaA, adjacent hexamers around the 3-fold symmetry axis were fitted using Segger tool in ChimeraX^60^. Finally, symmetry copies of SlaA and SlaB were generated using ChimeraX^61^. A combination of manual and automated rigid body fitting was employed only for domains in U4 due to low resolution.

---

**Supplementary Figure 1.**
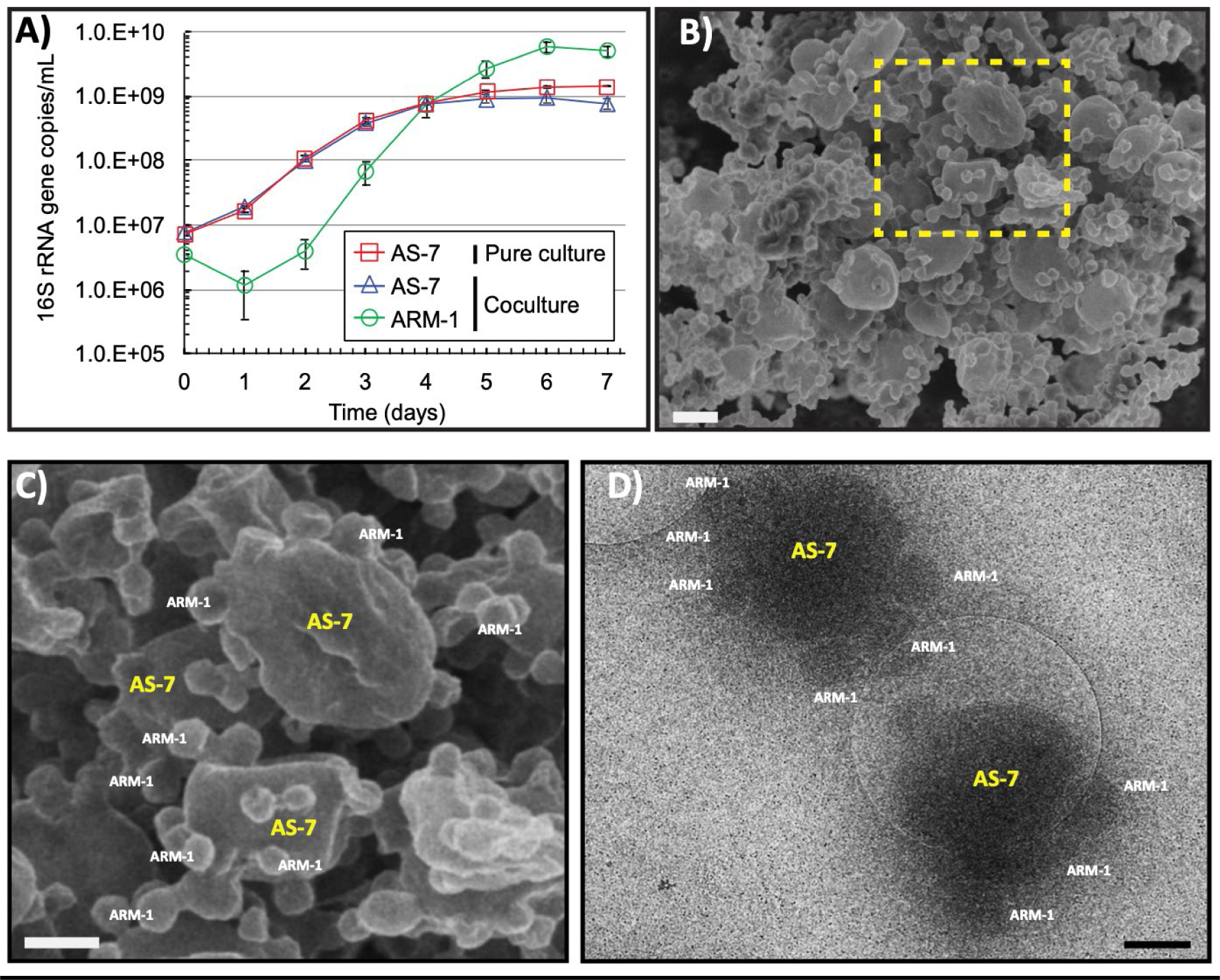
The ARM-1 and AS-7 co-culture, growth, and association. (A) Growth curve analysis of pure AS-7 and in co-culture with ARM-1 using 16S-rRNA quantification. (B) SEM analysis of an example co-culture used in this work showing small ARM-1 cells attached to larger AS-7 cells. (C) Zoomed in view of the image shown in (B) (yellow dotted square area) showing multiple ARM-1 cells are attached to each AS-7 cell. (D) Cryo-TEM image of ARM-1 and AS-7 co-culture showing ARM-1 and AS-7 association. Scale bar 500 nm.

**Supplementary Figure 2.**
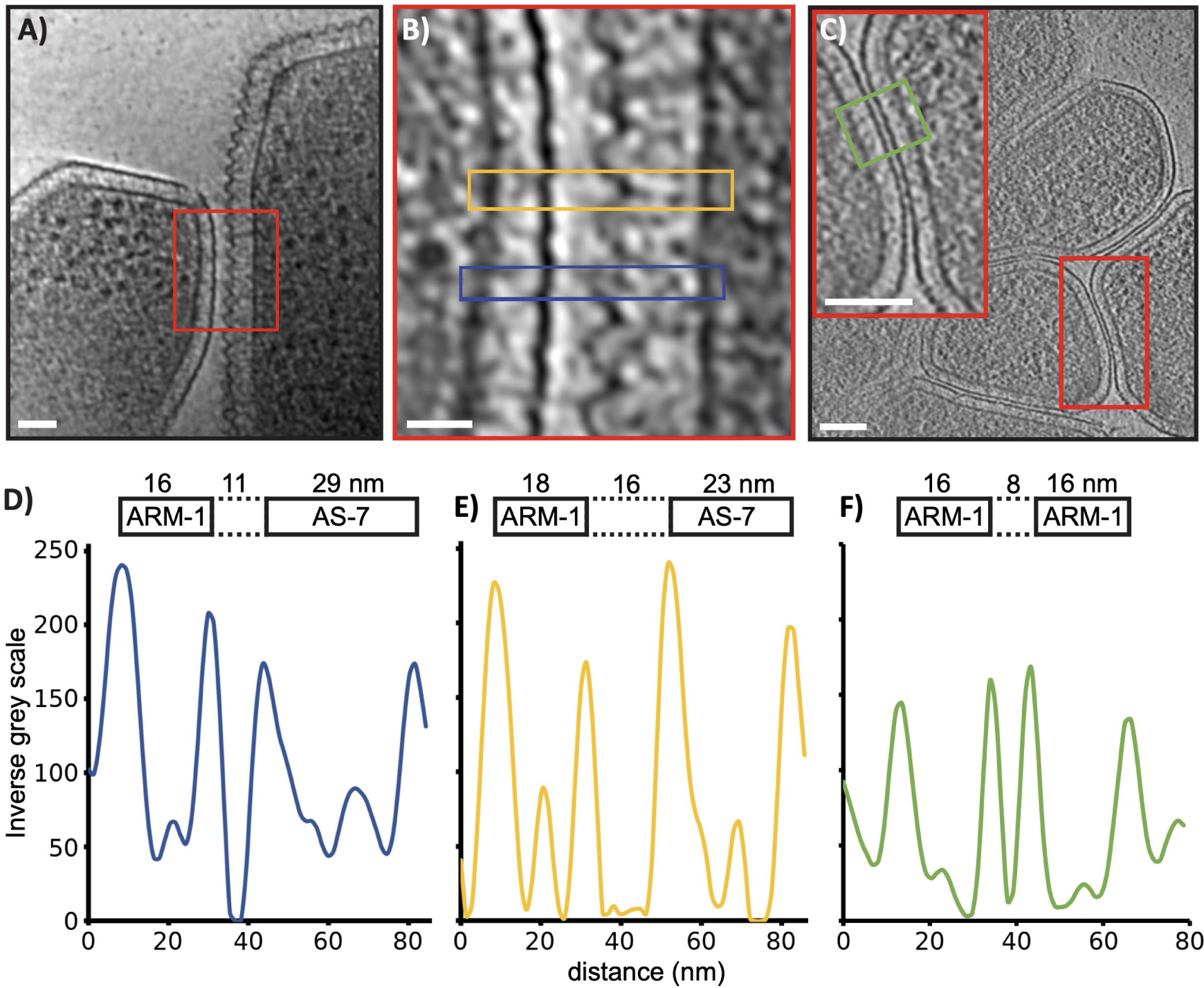
Interaction between ARM-1 and AS-7. (A) Tomographic slice of AS- 7 and ARM-1 co-culture showing an ARM-1 cell (left) interacting with an AS-7 host cell (right). (B) The red box in (A) is enlarged in (B). High-resolution details of the interface between ARM- 1 and AS-7 is visible. The AS-7 surface forms an undulating contour. The distance between ARM- 1 and AS-7 varies in peak (blue box) and trough (yellow) regions. (C) Tomographic slice showing interaction between two ARM-1 cells. Inset, zoomed in view of the region within red box showing the distance between two neighbouring ARM-1 cells is consistent along the length of their contact. (D-E) Density profile analyses of tomographic slices showing distance between an ARM-1 and a AS-7 cell at peak (D) and trough (E) regions and between two ARM-1 cells (F). Scale bar 20 nm (A), 20 nm (B), and 50 nm (C) respectively.

**Supplementary Figure 3.**
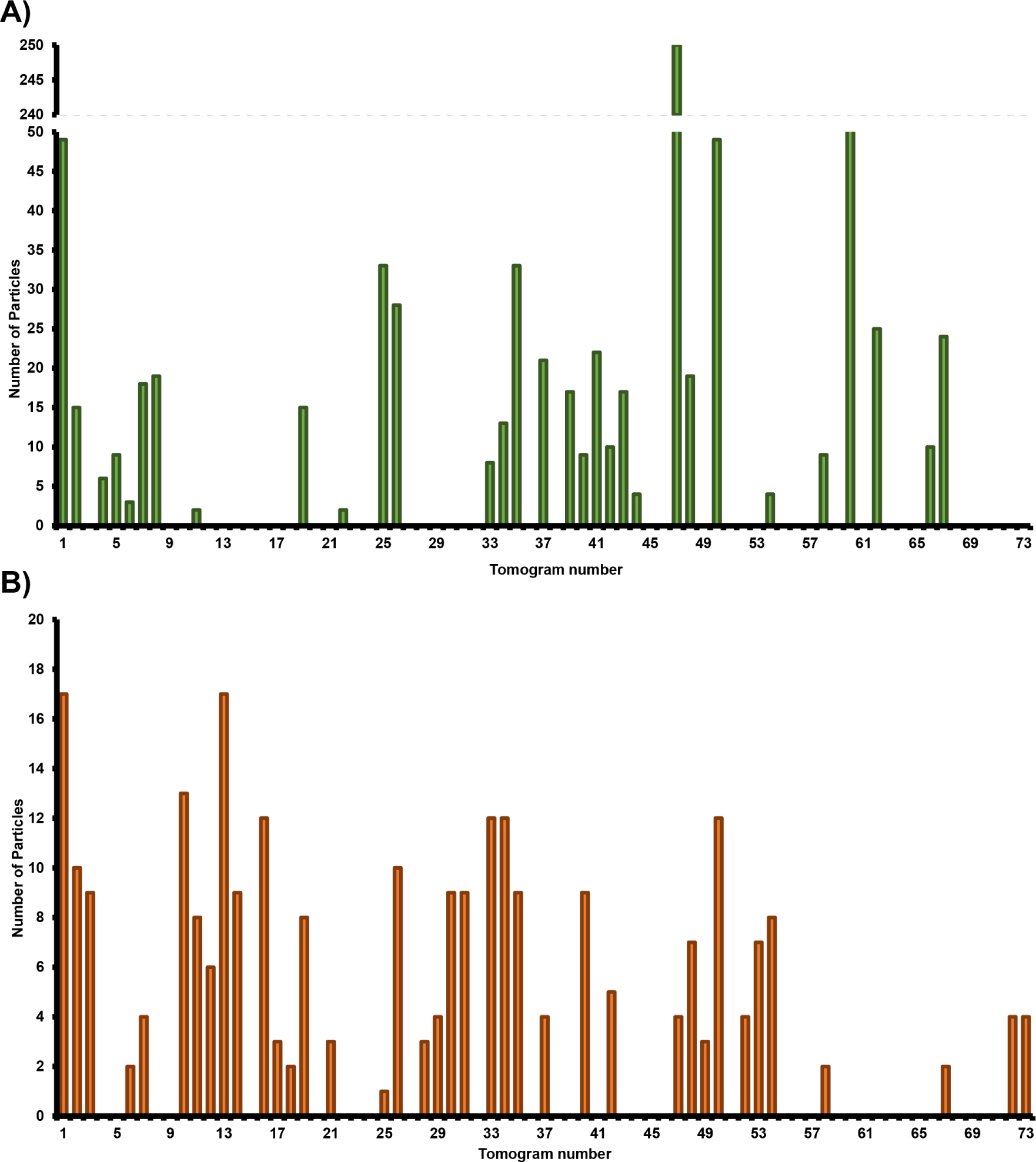
Number, frequency and distribution of particles picked from tomograms. **(A)** Number of “top-views” of primed tubes picked from each tomogram. **(B)** Number of extended tubes picked from each tomogram.

**Supplementary Figure 4.**
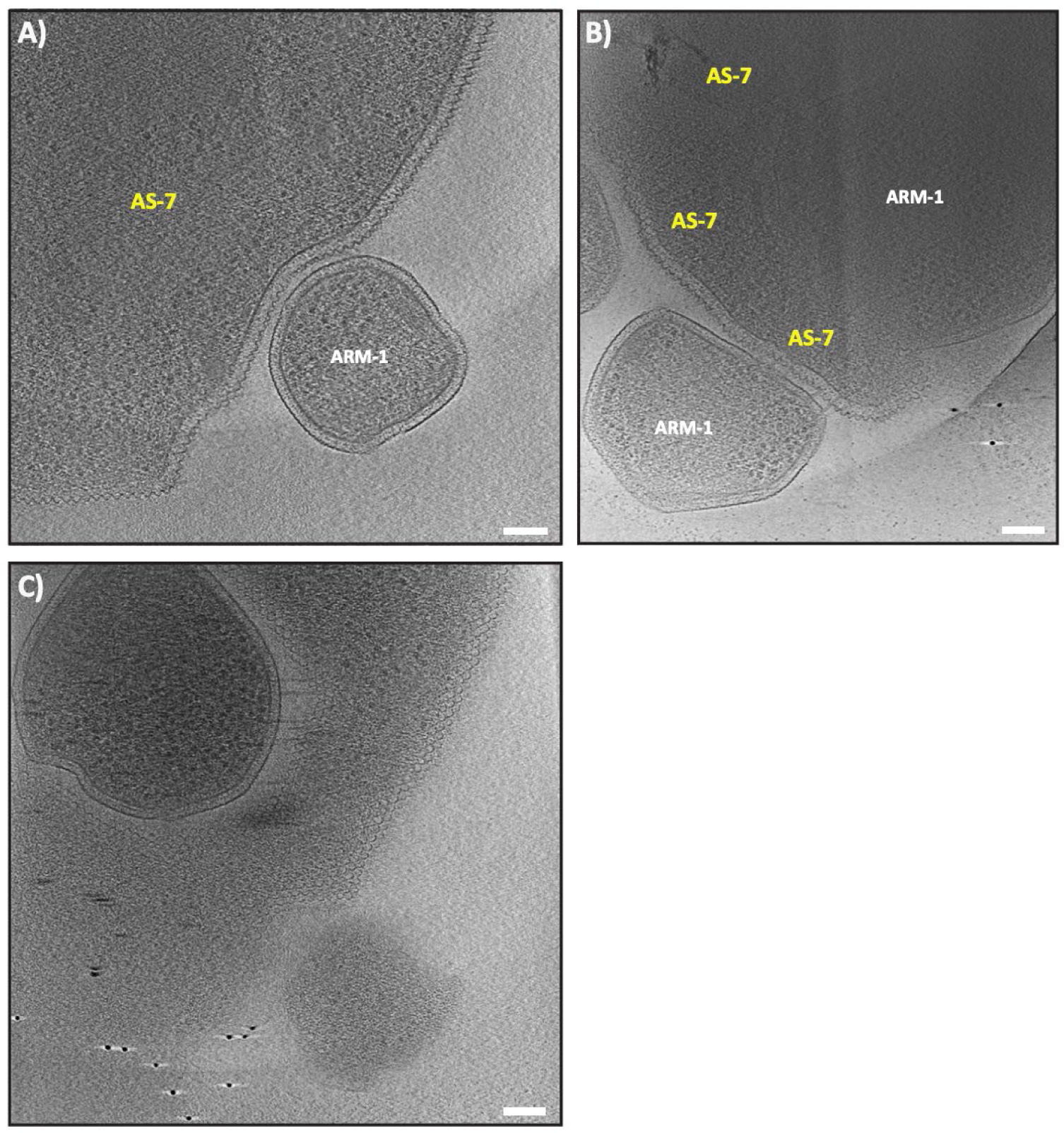
Potential engulfment of ARM-1 by AS-7. Tomographic slices of ARM- 1 and AS-7 coculture showing AS-7 forming a ‘phagocytic cup’-like structure around ARM-1 (A). An advanced stage of this the process where an ARM-1 cell is almost fully surrounded by a ‘pseudopodia-like’ extension from AS-7 (B). Finally, a tomographic slice showing a nearly internalised ARM-1 cell. SI Movie 3 provides a 3D view of (C). Scale bar 100 nm (A-C).

**Supplementary Figure 5.**
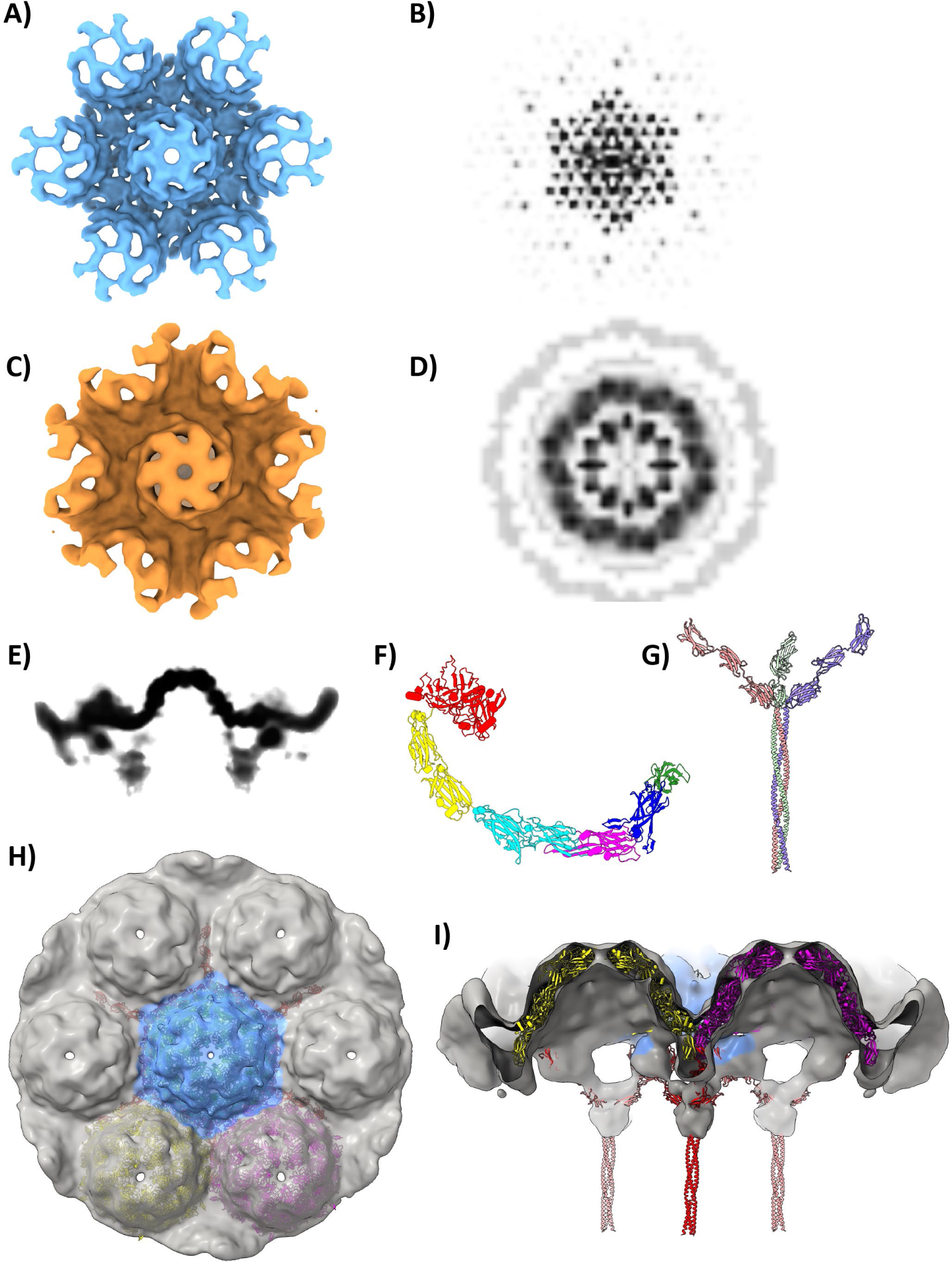
In situ structures of the AS-7 S-layer, primed nanotube and their respective power spectrums. (A) Top view of the surface representation of the AS-7 S-layer structure determined by STA. (B) Central cross section of the 3D power spectrum for (A) revealing clear six-fold symmetry. (C) Top view of the surface representation of the primed AS-7 nanotube. (D) Central cross section of the 3D power spectrum for (C). (E) Orthoplane view of the AS-7 S-layer structure as determined by STA. (F) AlphaFold model of SlaA^55^. Distinct domains marked in different colours - red (28-373), yellow (74-506, 507-639), cyan (640-784, 785-919), magenta (920-1076), blue (1077-1230) and green (1231-1345). Full-length model of SlaA was split into multiple units and fitted into the STA density map in a semi-automated manner ^56,58,59^. (G) AlphaFold multimer model of SlaB trimer^55^. Individual protomer are shown in red, blue and green. (H) Fits of the three representative hexamer models of SlaA shown in blue, yellow and magenta as ribbon representation within the transparent density map^57,60,61^. (I) Same as (H) rotated 90° about X-axis with cross section showing 4 SlaA proteins in yellow and magenta as well as SlaB trimers in the background.

**Supplementary Figure 6.**
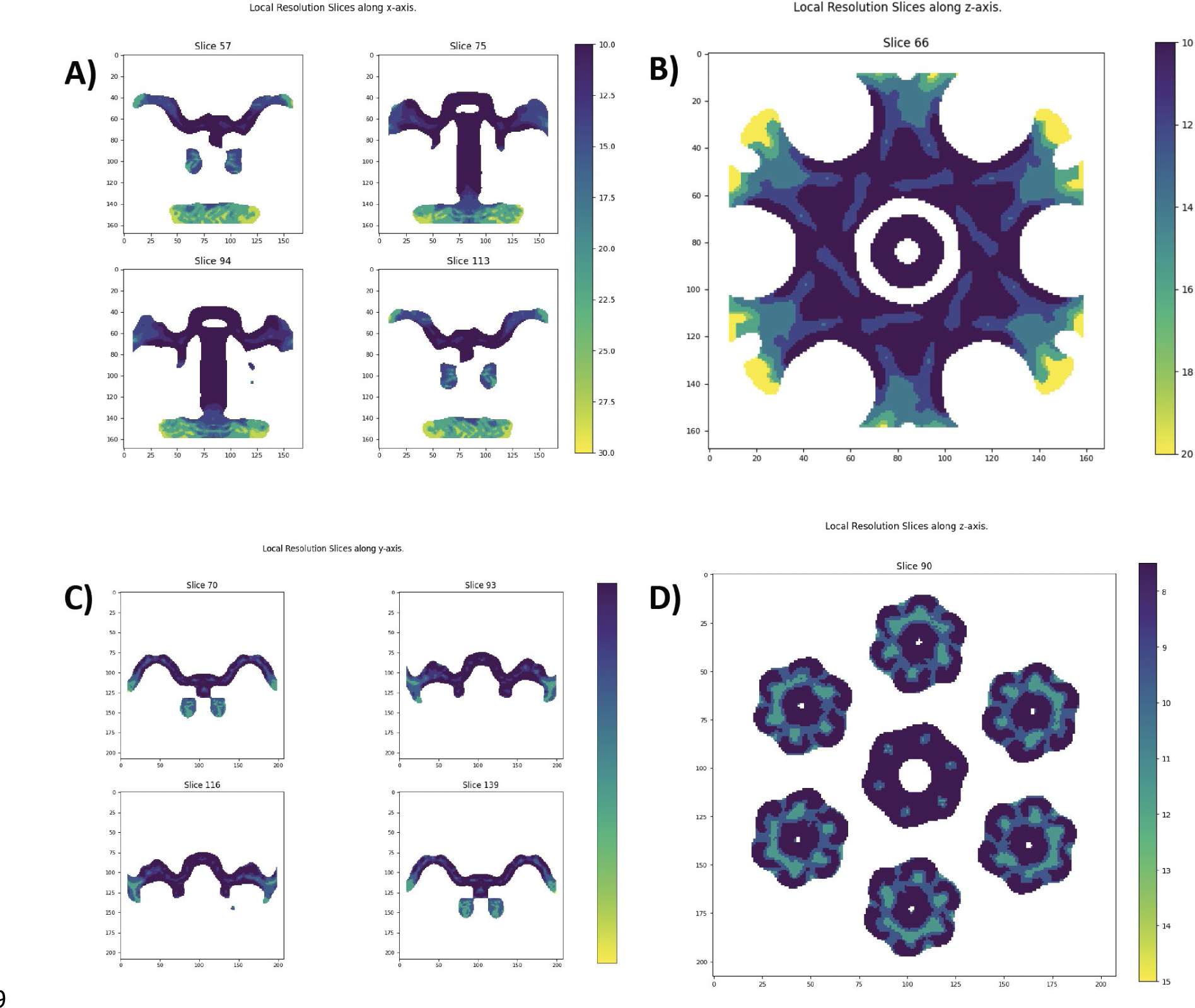
Local resolution of the subtomogram averages of AS-7 S-layer and AS-7 tube calculated by ResMap. (A) ResMap analysis of AS-7 tube structure, side cross-sections and top cross-section (B). (C) ResMap analysis of the AS-7 S-layer structure side cross-sections and top cross-section (D).

**Supplementary Figure 7.**
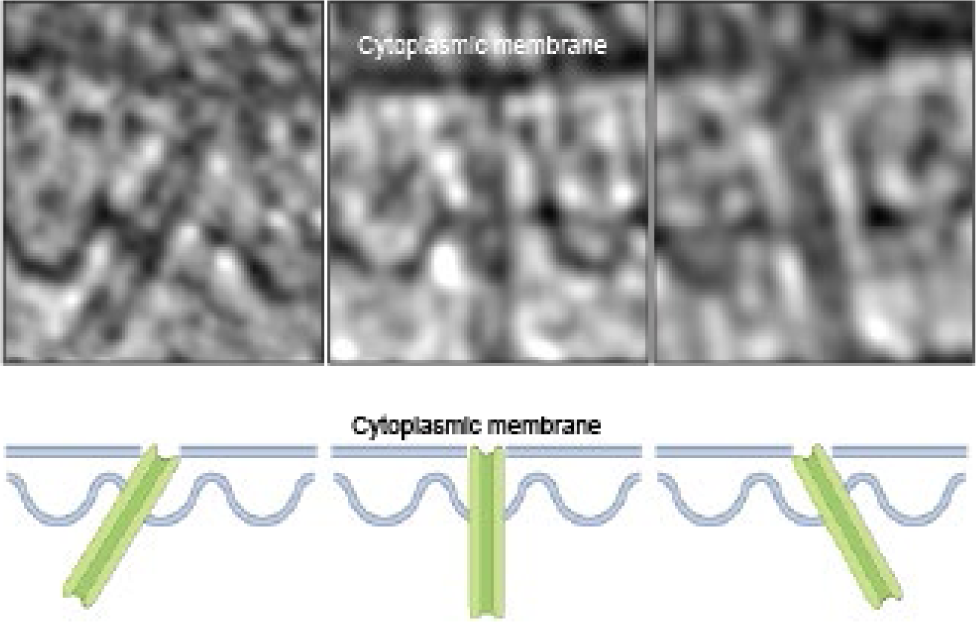
Three example particles of extended tube structures demonstrating the variability of their orientation relative to the AS-7 membranes (top). Diagrammatic representations of the tube orientations from the above panels. (bottom).

**Supplementary Figure 8.**
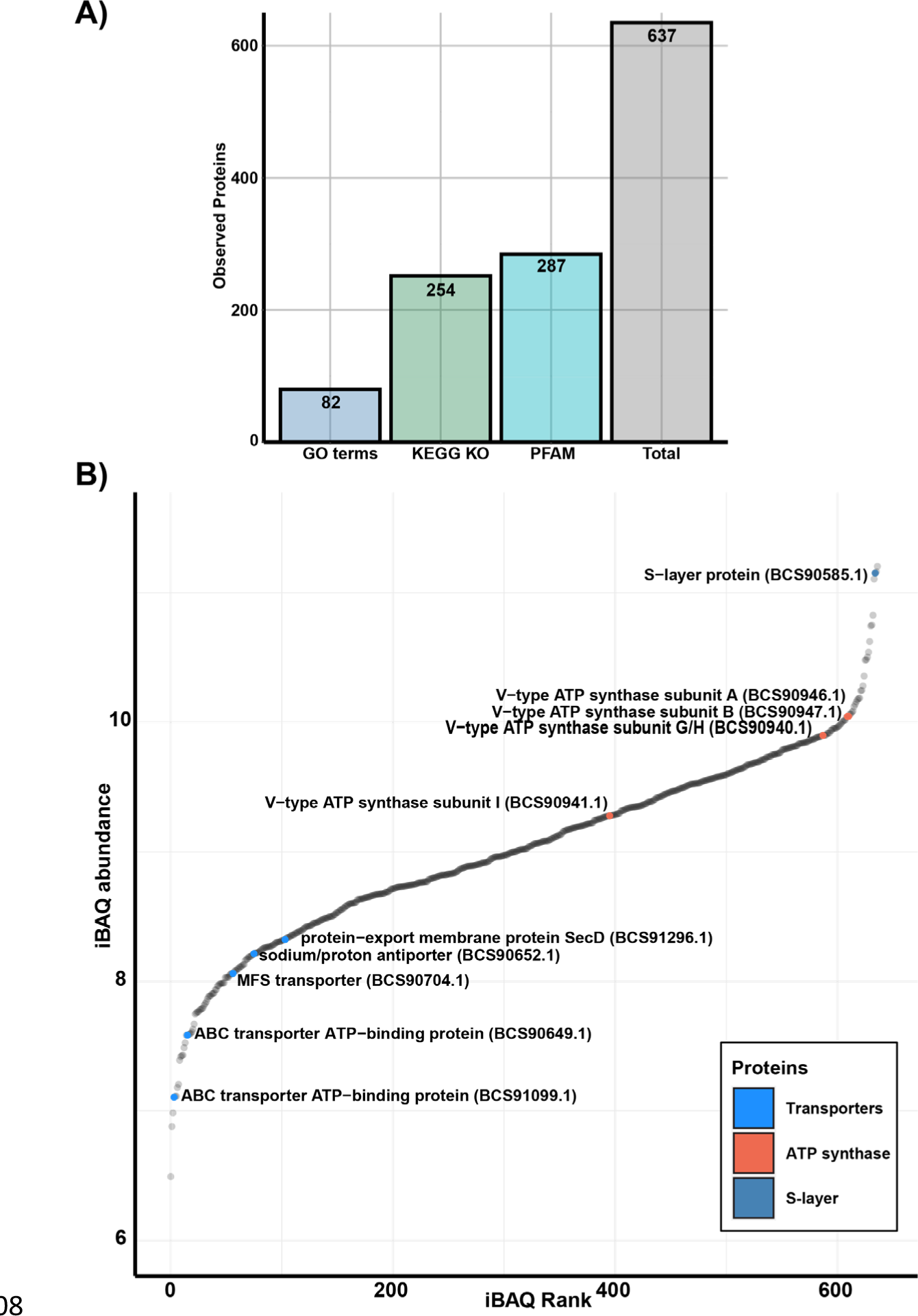
Proteomics analysis of ARM-1 during co-culture with AS-7. **(A)** Bar chart of automated annotations of ARM-1 proteins by GO term, KEGG KO, and PFAM. **(B)** ARM-1 protein abundance determined by iBAQ analysis, ATP synthases and transporters are highlighted in red and blue respectively. The ARM-1 S-layer is highlighted in dark blue.

**Supplementary Figure 9.**
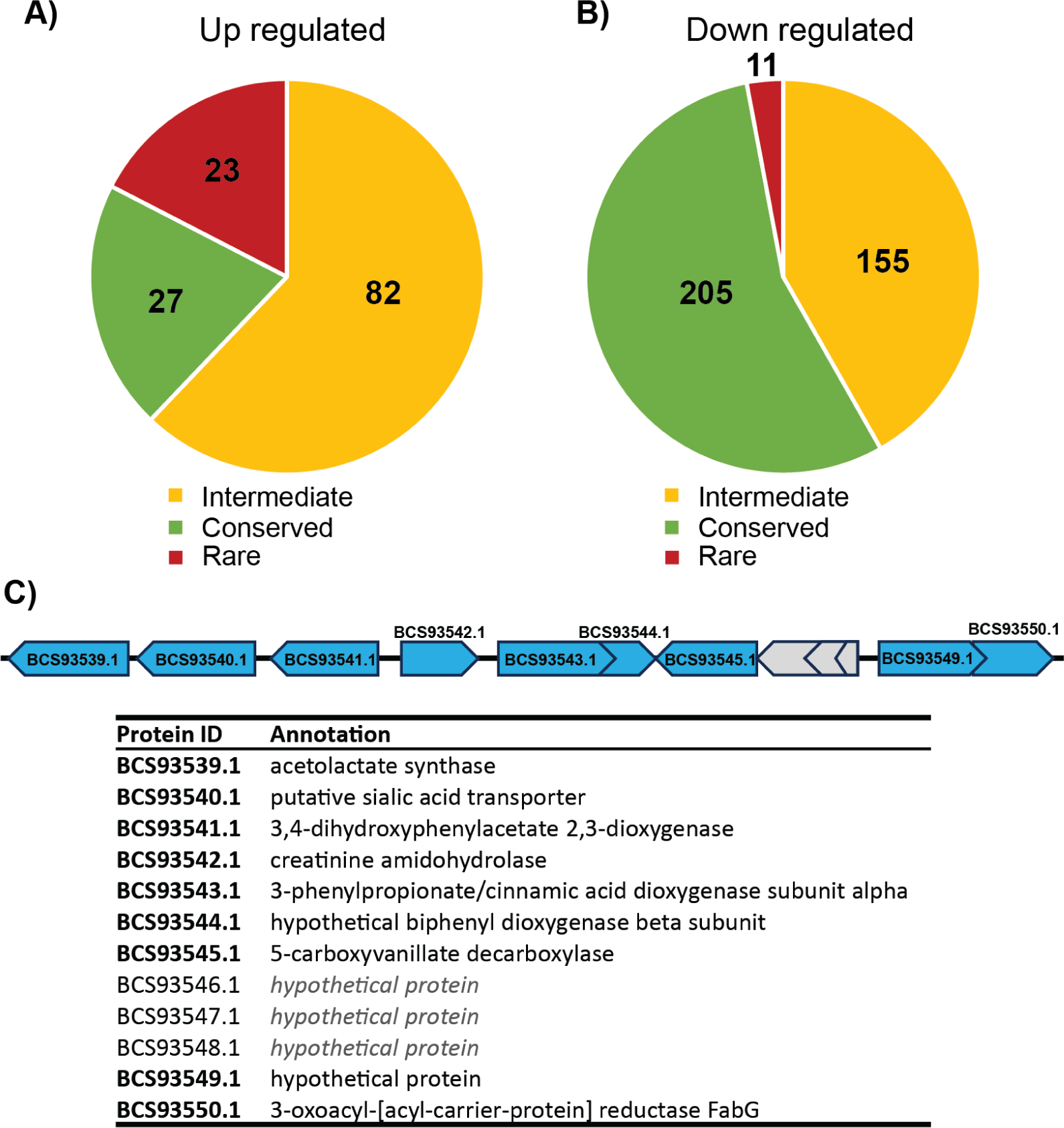
Comparative genomics analysis of Sulfolobaceae. Genes encoding proteins that were found to differentially regulated by proteomics analysis in this study were compared across the Sulfolobaceae family by presence absence analysis. A complete list of genomes used is in supplementary table 2. Pie charts show the level of conservation of all up regulated (**A**) and down regulated (**B**) proteins. Conserved genes (green) are those found in >80% of Sulfolobaceae genomes, Intermediate genes (yellow) are present in 10-80%, and rare genes (red) are present in <10% of Sulfolobaceae. (**C**) Gene cluster (ii), which is predicted to be involved in catabolism of aromatic compounds and is found in *M. javensis* but not in other Sulfolobaceae representatives. Genes with upregulated protein products in the presence of ARM-1 are shown in blue, while others are shown in grey. Automated annotations for each protein ID are shown below; these were generated by Prokka^62^.

**Supplementary Figure 10.**
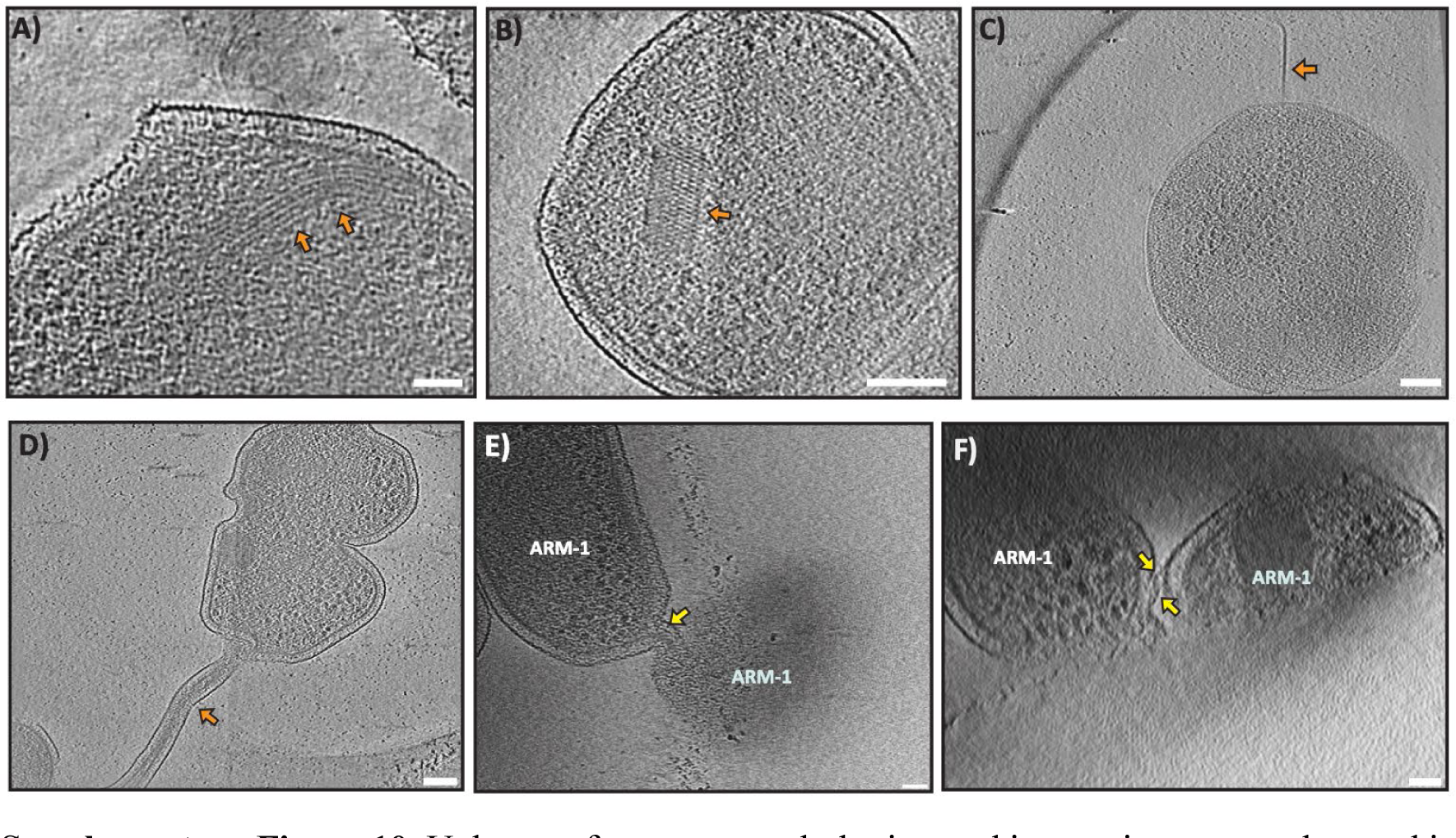
Unknown features, morphologies, and interaction events observed in tomograms of ARM-1 cells. (A) Tomographic slice of an ARM-1 cell with orange arrows highlighting unknown filamentous structures in the cytoplasm, measuring 15-20 nm in diameter. (B) Tomographic slice of an ARM-1 cell with an orange arrow pointing to an unknown trapezoidal-shaped density observed in the cytoplasm. This density is seen in several ARM-1 tomographic reconstructions and, while its length and width appear irregular, it consistently presents a cross-hatched appearance. (C) Tomographic slice of ARM-1 with an orange arrow indicating a density corresponding to an archaellum. (D) Tomographic slice of an ARM-1 cell showing a tube-like extension of the S-layer and cytoplasm, as highlighted by the orange arrow. (E-F) Intercellular tubes connecting cytoplasms of two DPANN cells.

## Supplementary Information Movie legends

**SI Movie 1**

Three-dimensional (segmented) view of an ARM-1 and AS-7 interaction showing intercellular proteinaceous tubes traversing between the host and ARM-1.

**SI Movie 2**

Three-dimensional (segmented) view of an ARM-1 and AS-7 interaction showing intercellular proteinaceous tubes traversing between the host and ARM-1 and unknown structural features inside ARM-1 cells.

**SI Movie 3**

Tomogram movie moving through the z axis showing the engulfment of an ARM-1 cell by the AS-7 host cell membrane.

**SI Movie 4**

Pseudo-atomic model of the AS-7 S-layer, showing the fit of the SlaA and SlaB AlphaFold models into the *in situ* AS-7 S-layer subtomogram average.

**SI Movie 5**

Structure of the contracted tube after segmentation, showing the cross-section of the tube from the perspective of inside the AS-7 cell and side on.

**SI Movie 6**

Tomogram movie through the z axis of an AS-7 cell, showing the presence of tube structures in the host cell in a pure host culture.

**SI Movie 7**

Tomogram movie moving through the z axis showing the membrane nanotube connections between ARM-1 cells.

## Supplementary Tables

**Supplementary table 1**

Proteomics analysis of AS-7 pure culture and AS-7 ARM-1 coculture.

**Supplementary table 2**

Presence loss analysis by comparative genomics analysis of Sulfolobaceae

## Notes

### Competing Interest Statement

The authors have declared no competing interest.

